# Semi-supervised single-cell cross-modality translation using Polarbear

**DOI:** 10.1101/2021.11.18.467517

**Authors:** Ran Zhang, Laetitia Meng-Papaxanthos, Jean-Philippe Vert, William Stafford Noble

**Affiliations:** Department of Genome Sciences, University of Washington; Google Research, Brain Team; Paul G. Allen School of Computer Science and Engineering, University of Washington

## Abstract

The emergence of single-cell co-assays enables us to learn to translate between single-cell modalities, potentially offering valuable insights from datasets where only one modality is available. However, the sparsity of single-cell measurements and the limited number of cells measured in typical co-assay datasets impedes the power of cross-modality translation. Here, we propose Polarbear, a semi-supervised translation framework to predict cross-modality profiles that is trained using a combination of co-assay data and traditional “single-assay” data. Polarbear uses single-assay and co-assay data to train an autoencoder for each modality and then uses just the co-assay data to train a translator between the embedded representations learned by the autoencoders. With this approach, Polarbear is able to translate between modalities with improved accuracy relative to state-of-the-art translation techniques. As an added benefit of the training procedure, we show that Polarbear also produces a matching of cells across modalities.

## 1 Introduction

Single-cell measurements are immensely valuable for quantifying the variance of certain forms of molecular activity within the cell and for identifying and characterizing cell subpopulations within complex tissues. However, a weakness of most single-cell assays is that only a single form of activity can be measured for each cell.

Consequently, a variety of machine learning methods have been proposed to translate between different types of single-cell measurements [1–6]. Typically, co-assay data is used to train such models in a fully supervised fashion. However, in practice co-assay measurements are often more challenging to produce and lower throughput than standard single-cell measurements. Furthermore, co-assays have only been developed relatively recently and thus are less abundant than single-assay data. Although the recently released Cobolt [7] and MultiVI [6] methods incorporate single-assay data into model training, neither directly assessed whether adding single-assay data from unrelated public datasets improves cross-modality translation. We hypothesize that training a translation model using both labeled data (co-assay data) and unlabeled data (single-assay data) from unrelated studies will result in more accurate translation performance than training from co-assay data alone.

Here we propose Polarbear, a semi-supervised approach that learns to translate between single-cell modalities by leveraging both single-assay and co-assay data. We focus on translation between scRNA-seq and scATAC-seq; i.e., given a scRNA-seq profile of a cell, Polarbear will produce as output the scATAC-seq profile of that same cell, and vice-versa. Polarbear is applicable to several types of co-assays that measure expression and chromatin accessibility within single cells, including CAR-seq [8], SNARE-seq [9], and Paired-seq [10]. Polarbear operates in two phases (Figure 1). In the first phase, we train two deep variational autoencoder (VAE) neural networks that learn, in an unsupervised fashion, to reduce each given type of data to a latent representation (the encoder) and then expand that representation to recover the original data (the decoder). (V)AEs have already been successfully applied to scRNA-seq and scATAC-seq data, primarily for the purpose of de-noising [11–17]. Polarbear trains one VAE for each type of data, while taking into consideration sequencing depth and batch factors [15, 16]. In phase two, we stitch together the encoder for one data type with the decoder of a second data type, interposing between them a single, fully connected “translator” layer. During this phase, the parameters of the encoder and decoder are frozen, and the translator parameters are trained in a supervised fashion using co-assay data. Repeating this procedure in reverse, Polarbear allows for bidirectional translation between scRNA-seq and scATAC-seq data. In principle, our method can also be applied to co-assays operating on other data modalities.

**Figure 1:**
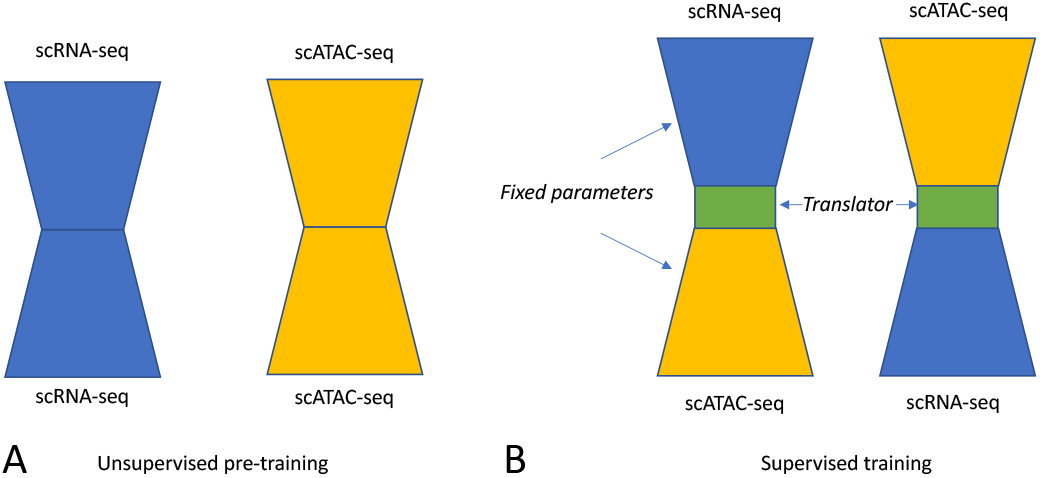
Polarbear’s semi-supervised framework. **A** In stage 1, Polarbear trains an autoencoder for each data modality, using both single-assay and co-assay data. **B** In stage 2, the encoder from one modality is stitched together with a decoder from the other modality (and vice versa), and the translation layers are trained in a supervised fashion using co-assay data.

In order to evaluate the performance of Polarbear, we propose a set of evaluation metrics for single-cell translation tasks, with the aim of teasing out several important aspects of translation performance. A drawback of current methods lies in the choice of evaluation metrics used for cross-modality profile prediction. Many previous methods report the correlation (for scRNA-seq) or area under the receiver operating characteristic curve (AUROC, for scATAC-seq) between the overall observed and predicted profiles. However, these performance measures can be strongly driven by the average profiles across cells, failing to reflect whether the prediction method accurately captures cell-to-cell variation. While MultiVI systematically demonstrates that the proposed method can predict differential expression between cell clusters or cell types, it does not address whether the model accurately captures differences among single cells.

Using our expanded set of performance measures, we demonstrate that Polarbear’s translation performance improves when we add single-assay data to the training procedure in the first phase. We also show that Polarbear outperforms BABEL [2], a state-of-the-art translation method, using several different performance measures. Finally, we demonstrate that Polarbear can be used to accurately match cells between modalities. Overall, our work illustrates the utility of exploiting single-assay data to aid in the prediction of cross-modality profiles.

### 1.1 Related work

Several previous methods have been developed for cross-modality prediction (Table 1). TotalVI builds a VAE that takes as input the concatenation of gene and protein expression profiles from the CITE-seq co-assay. The autoencoder learns to impute missing protein expression profiles based on scRNA-seq profiles [1]. BABEL translates between single-cell modalities by joining two autoencoders, one from each data domain [2]. The scMM method uses a mixture-of-experts multimodal deep generative model to learn a joint embedding between modalities and predict missing modalities [3]. Seurat predicts the missing domain profile of a cell by identifying neighboring cells in co-assay data and then computing the average profile of those neighbor cells in the second modality [4]. Multigrate jointly embeds data from two or more modalities and uses the joint embedding to infer profiles in each domain [5]. MultiVI embeds scRNA-seq and scATAC-seq profiles into a shared space by joining two VAEs. The trained model is then able to take single-assay data as input and predict the missing modality [6].

**Table 1:**
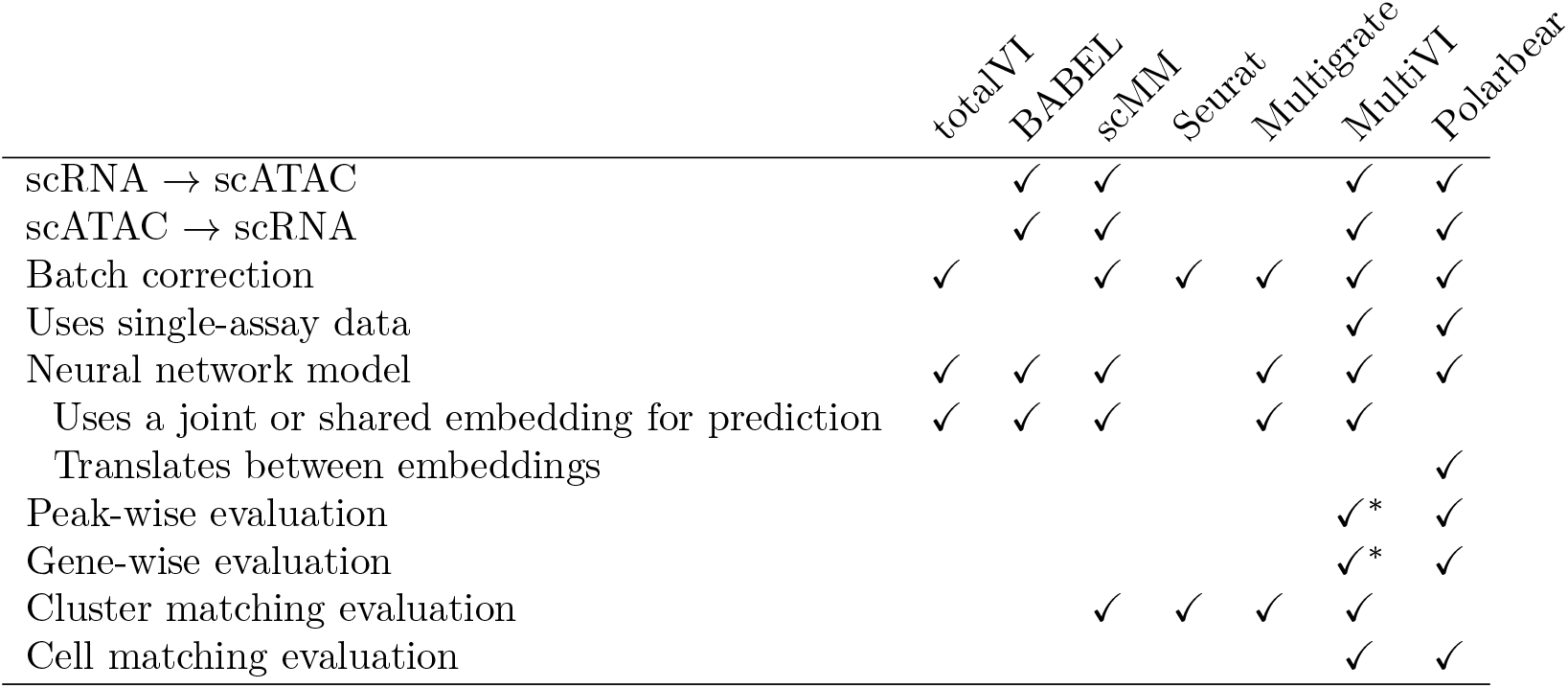
Method comparison. *Evaluation is done across cell types per-peak or per-gene, rather than across individual cells.

Polarbear differs from previous translation models in several ways. First, Polarbear uses a stepwise optimization approach to first learn embeddings with both single and co-assay data and then translate between embeddings across modalities based on co-assays, whereas other methods optimize the entire model jointly with a weighted sum of losses for each task. The separate optimization steps make Polarbear less likely to be biased toward optimization of a specific task and requires less hyperparameter tuning. Second, previous models generate predictions based on a joint or shared embedding of both modalities. Polarbear does not require a shared or joint embedding; instead, it adds a translation layer between the embeddings across modalities based on co-assay data. Thus Polarbear is more flexible at leveraging single-assay data and incorporating pre-trained models from new data modalities.

More importantly, Polarbear is able to use single-assay data collected from public datasets to improve its translation performance. Although MultiVI can learn from single-assay data, the question of whether adding single-assay data from unrelated datasets improves translation performance has not been addressed.

In this study, we choose to compare our method with BABEL, for several reasons. First, BABEL directly addresses the task of translating between scRNA-seq and scATAC-seq, and it has been applied to the SNARE-seq co-assay data we use in this study, so it is most likely we can make a fair comparison with BABEL. We attempted to run multiVI, which was published on bioRxiv very recently, but it ran out of memory when trained on the SNARE-seq data. Second, because the focus and novelty of Polarbear is the semi-supervised framework that leverages single-assay data from unrelated studies, instead of comparing extensively with current methods that are not specifically designed for this task, we demonstrate the power of our semi-supervised framework leveraging single-assay data (“Polarbear”) by comparing it with a Polarbear model that is only trained on co-assay data (“Polarbear co-assay”). We foresee that Polarbear’s semi-supervised framework could be adapted to other existing architectures to boost their translation performance.

## 2 Methods

### 2.1 Polarbear model

Polarbear’s autoencoder model adopts ideas from scVI and peakVI [15, 16], which take into account sequencing depth and batch factors. Specifically, Polarbear has the following architecture (Figure 2). One of Polarbear’s encoders takes gene expression raw counts as input and assumes they follow zero-inflated negative binomial (ZINB) distributions. The other encoder takes in binarized scATAC counts and assumes they follow Bernoulli distributions. To save memory for the scATAC parts of the model, we only allow for within-chromosome connections in the first two encoder layers and the last two decoder layers. We first specify the latent dimension of each autoencoder, and then we define the dimension of each hidden layer as half of the geometric mean between the input and latent dimensions. To correct for batch effects, we one-hot encode batch factors and concatenate them to the input and embedding layers. To correct for sequencing depth differences across cells, Polarbear includes a sequencing-depth factor as the sum of counts per cell, which is used to calculate the ZINB loss (for scRNA-seq) and binary cross entropy loss (for scATAC-seq), together with other distributional variables learned from the embedding layers in the corresponding modality. The loss of each VAE is the sum of the reconstruction loss and a weighted KL divergence loss.

**Figure 2:**
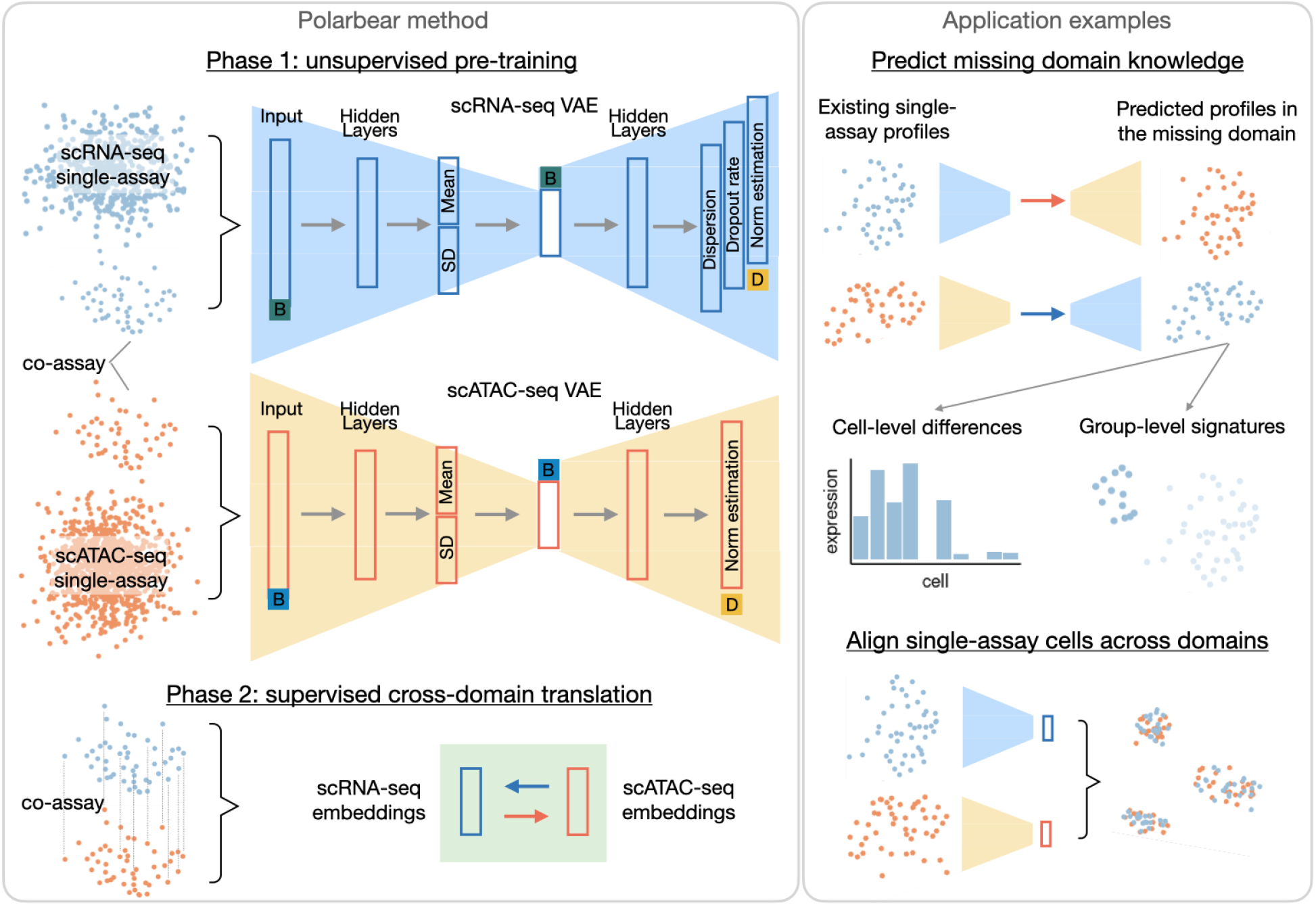
Polarbear’s semi-supervised framework and applications. Left: Polarbear’s semi-supervised framework. In Phase 1, Polarbear trains a variational autoencoder for each data modality, using both single-assay and co-assay data. In Phase 2, the encoder from one modality is stitched together with a decoder from the other modality (and vice versa), and the translation layers are trained in a supervised fashion using co-assay data. Specifically, the VAEs take into account sequencing depth (D) and batch effects (B). The scRNA-seq VAE assumes that counts are drawn from a zero-inflated negative binomial distribution, and the scATAC-seq VAE assumes a Bernoulli distribution. Right: Sampled applications of Polarbear. Polarbear can predict missing domain profiles based on the known domain, capturing individual cell-level differences and group-level signatures in the missing domain. Furthermore, Polarbear can match single cell profiles across modalities.

After the autoencoders in both domains are optimized, we learn a single translation layer between the embedding layers of the scRNA-seq and scATAC-seq, supervised by co-assay data, to minimize the translation loss on each modality.

In both translation directions, since the distributional variables are independent of sequencing depth, the size-normalized expectations of the distributions (i.e. the “norm estimation” in Figure 2) can be directly used for subsequent tasks such as differential expression analysis. Because the true scATAC-seq profiles used for evaluation are unnormalized binarized counts, we further generate an “unnormalized” prediction so that we can evaluate on the true profiles and make a fair comparison to methods that do not take into account sequencing depth. Since there is a clear correlation between the scATAC-seq and scRNA-seq depth factor when both modalities are observed, which may capture both technical (e.g. batch effect) and biological (e.g. cell cycle) effects, we predict the scATAC-seq depth factor in the translation task based on the known scRNA-seq profile. Specifically, we use ridge regression for this prediction task, with a penalty term determined by cross-validation within the training set. Finally, we multiply the learned sequencing depth with the normalized predictions to generate unnormalized predictions in the test set.

### 2.2 Hyperparameter tuning

The Polarbear neural network architecture has two primary hyperparameters: the latent dimensions of the autoencoders, and the weight of the KL divergence term in each VAE. In this study, we use the validation set to choose hyperparameters, selecting the number of latent dimensions ({10, 25, 50}) and the KL divergence weight ({0, 0.5, 1, 2}). In the random test set setup, we randomly split the SNARE-seq dataset, assigning 60% of cells to the training set, 20% to the validation set, and 20% to the test set. In the unseen cell type scenario, we use the same validation and test set as BABEL, where the validation and test set are the largest two cell clusters based on the SNARE-seq scRNA-seq dataset. The rest of cells are used as the training set.

We downloaded BABEL’s scripts and followed the instructions to generate predictions on SNARE-seq [2]. We verified that we are able to reproduce the performance reported in the paper. In all scenarios, we make sure that BABEL’s train/validation/test splits are the same as Polarbear’s. BABEL has proposed a set of default parameters (latent dimension: 16, weight factor: 1.13); however, for a fair comparison we tune the following 2D grid of hyperparameters: number of latent dimensions in {10, 16, 25, 50} and weight factor to balance scATAC loss in {0.2, 1.33, 5}. We then select BABEL’s best performing model based on each task’s performance on the validation set.

### 2.3 Performance measures

In designing performance measures for cross-modality translation, we tried to place ourselves in the shoes of a prospective end user of our predictive model. Imagine a scenario in which we are interested in leveraging an existing scRNA-seq dataset to predict chromatin accessibility in a particular biological system, applying our trained model to the scRNA-seq matrix to yield a predicted matrix of scATAC values. Given the predicted peak activations, we can imagine trying to solve two different problems.

In the first setting, we begin by identifying cell types using the original scRNA-seq data or identifying cell groups based on the experimental design (e.g. disease and control groups). We may then be interested in the pattern of predicted chromatin peak activations within each cell type or group. In this setting, a classifier-based measure such as AUPR or AUROC, computed separately for each peak, would accurately capture the per-peak predictive behavior across single cells and thus be indicative of per-peak predictive power across clusters or groups. Each of these two measures has advantages. The AUPR emphasizes enriching the top of the ranked list of predictions with positives. On the other hand, AUROC explicitly corrects for differences in “skew” (i.e., differences in the number of non-zero values) for each peak. To correct for the skew in AUPR measurement toward peaks with large positive proportions (PP defined as #(cells with peak expressed)/#cells), we calculate AUPRnorm = (AUPR-PP)/(1-PP), where 0 represents the behavior or a random predictor and 1 indicates perfect predictor. In this work, we use AUROC and AUPRnorm as performance measurements.

In the second setting, we can use the profile of predicted peak activations across each cell to match scRNA-seq profiles to corresponding scATAC-seq profiles. In this setting, we identify these matches based on Euclidean distance. We want to ensure that each predicted profile’s nearest neighbor is the correct match; hence, we can use the fraction of samples closer than the true match (FOSCTTM) as a performance measure [18].

Based on these scenarios, we report here the average per-peak AUROC and AUPRnorm, as well as the FOSCTTM. Similarly, when predicting gene expression from scATAC-seq, we report the average per-gene Pearson correlation (on log-scaled expression) and the FOSCTTM. We do not foresee a scenario in which the overall “flattened” performance of the model, in which we treat all values in the matrix as a single list and compute a single score (Pearson correlation, MSE, AUPR, or AUROC), will be of primary interest to an end user.

### 2.4 Single-cell data pre-processing

For co-assay data, we use SNARE-seq data from mouse adult brains (~10k cells) [9]. We filter out peaks that occur in fewer than 5 cells or more than 10% of cells, and we filter out genes that are expressed in fewer than 200 cells or more than 2500 cells, as in the original SNARE-seq paper. To learn robust representations of each domain, we first train the autoencoders using the SNARE-seq data combined with publicly available scRNA-seq and scATAC-seq profiles from adult mouse brains [19–21]. We collected scRNA-seq profiles from ~160k cells and scATAC-seq profiles from ~855k cells, and we randomly downsampled the latter dataset to ~170k cells for use in training (Table 2).

**Table 2:** Data sets.

| Data set | Cells | Assay | Platform |
| --- | --- | --- | --- |
| SNARE-seq | ~10k | Co-assay | Illumina HiSeq 2500/4000 |
| Li *et al.* | ~800k | snATAC-seq | Illumina HiSeq 2500 |
| Fang *et al.* | ~55k | scATAC-seq | Illumina HiSeq 2500 |
| Zeisel *et al.* | ~160k | scRNA-seq | 10x Genomics |

For scRNA profiles, we use SNARE-seq genes as a reference, map genes in the other datasets to the gene symbols in the SNARE-seq data, and further filter out non-protein-coding genes based on Gencode annotations [22]. We remove sex chromosome genes for consistency across datasets. In this way, 17,271 genes are maintained for input to Polarbear. For the scATAC profile, we first lift all peaks to mm10 [23]. Because the ATAC-seq peak locations vary across datasets, we use the SNARE-seq peaks as a reference and map features from other datasets to SNARE-seq peaks if there is an overlap of 1 bp or more. Peaks from sex chromosomes are again filtered out. In the end, 220,526 peaks are input to the Polarbear model.

### 2.5 Cluster-level analysis

A common task in single-cell analysis is to cluster the cells according to the similarity of their scRNA-seq or scATAC-seq profiles; accordingly, a good translator should be able to produce predicted profiles that yield clusters similar to the clusters produced by the observed data.

An important task for our predicted profile is to be able to predict cell-type specific marker genes. To evaluate this, we use the clustering analysis performed in the original SNARE-seq study, which yielded an expert-curated list of marker genes for each cell cluster [9]. This analysis also identified genes and peaks that are significantly highly expressed in each cell type.

To calculate whether a gene or a peak is specifically expressed in a specific cell type, we label cells in the corresponding cell type as positive and cells in other cell types as negative. We then calculate the AUROC of the predicted gene/peak-wise profile relative to these labels. A high AUROC score suggests the gene/peak is specifically expressed in the corresponding cell type. To calculate differentially expressed genes for a cell type, we perform a one-sided Wilcoxon rank-sum test between the expression pattern in the corresponding cell type and that in unrelated cell types, and we control the FDR using the Benjamini–Hochberg procedure. To validate differential expression predictions, we label differentially expressed genes derived from the true profile (FDR ≤ 0.01) as positive and other genes as negative, and we calculate a precision-recall curve using the predicted differential expression p-value.

## 3 Results

### 3.1 Polarbear accurately translates between single-cell data domains

We begin by testing Polarbear’s ability to translate between scRNA and scATAC profiles in a SNARE-seq adult mouse brain co-assay dataset. To learn robust representations of each domain, we train the autoencoders with large-scale, publicly available scRNA and scATAC single-assay profiles, also derived from adult mouse brains (see Section 2.4). We then train Polarbear’s translator layer in a supervised fashion using a training set of 80% of the cells from the SNARE-seq dataset, evaluating translation performance on the test set comprised of the remaining 20%.

To evaluate the performance of our model, we measure how well the predicted scRNA-seq profiles allow us to recapitulate gene expression differences across cells. To do this, we calculate the gene-wise correlation between the predicted profile and the true normalized profile [24]. In this analysis, Polarbear outperforms BABEL, yielding an improved correlation for 1067 out of 1205 genes (Wilcoxon rank-sum test p-value 4.51 × 10^−33^, Figure 3A, Table 3). We also observe that Polarbear strongly outperforms a Polarbear variant (“Polarbear co-assay”) that is trained only with co-assay data (Figure 3B). For translation in the opposite direction (scRNA-seq → scATAC-seq), we calculate the peak-wise area under the receiver operating characteristic curve (AUROC) of the predicted profile relative to the observed, binarized scATAC-seq profile. Polarbear outperforms both competing methods in this scenario (Figure 3C–D, Supplementary Figure 1A–B, Table 3).

**Table 3:** **Translation performance** represented by median and interquartile range (IQR).

| test set | evaluation metric | BABEL | Polarbear co-assay | Polarbear |
| --- | --- | --- | --- | --- |
|  |  | median (IQR) | median (IQR) | median (IQR) |
| random 20% | gene-wise correlation | 0.131 (0.141) | 0.146 (0.147) | 0.195 (0.185) |
|  | peak-wise AUROC | 0.636 (0.0872) | 0.647 (0.0839) | 0.663 (0.0845) |
|  | peak-wise AUPRnorm | 0.0136 (0.0153) | 0.0156 (0.0167) | 0.0190 (0.0199) |
| unseen cell type | gene-wise correlation | 0.0701 (0.0667) | 0.0561 (0.0985) | 0.122 (0.0999) |
|  | peak-wise AUROC | 0.560 (0.0871) | 0.564 (0.0784) | 0.592 (0.0876) |
|  | peak-wise AUPRnorm | 0.00795 (0.0128) | 0.00896 (0.0133) | 0.0132 (0.0183) |

**Figure 3:**
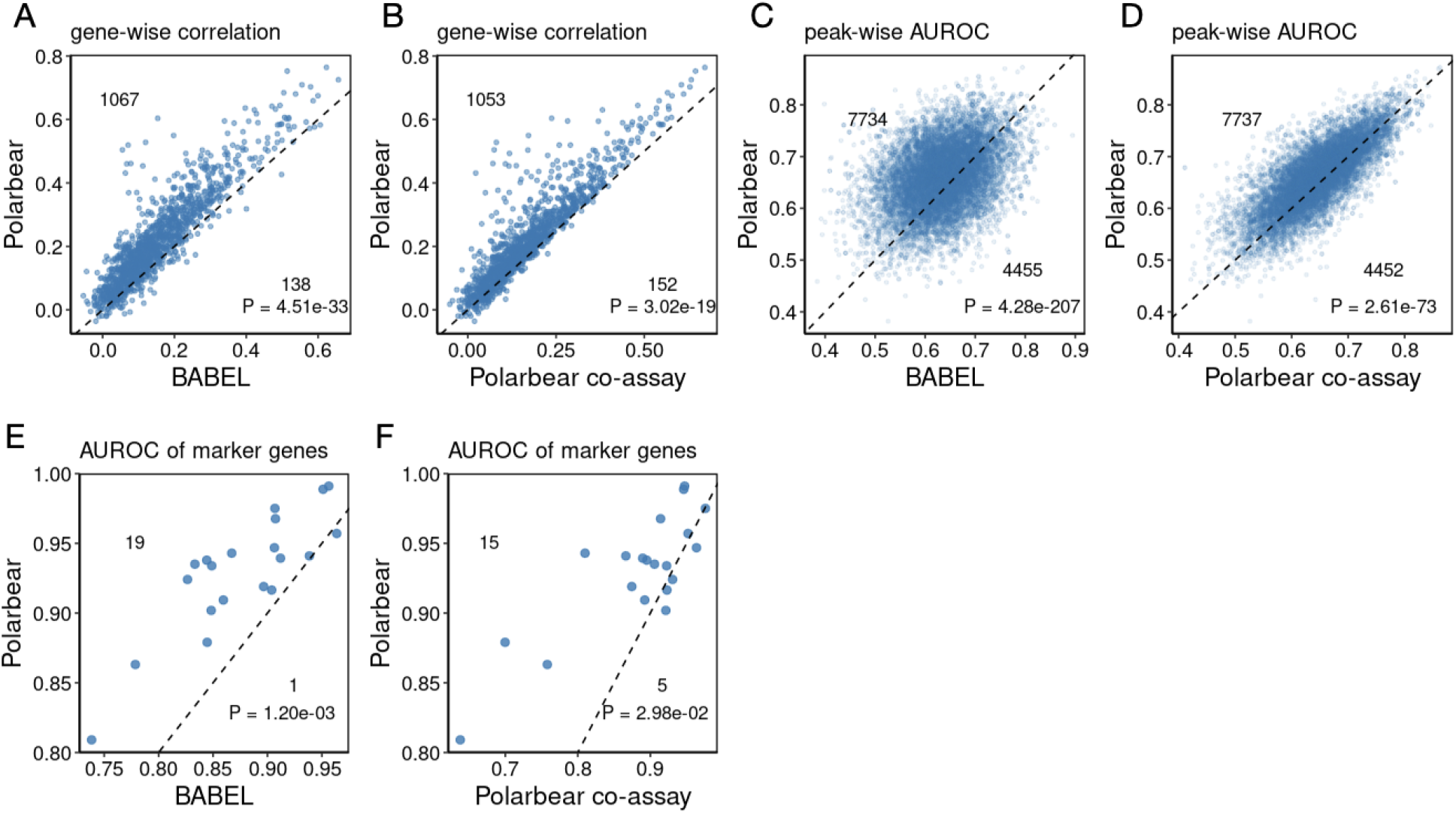
Cross-modality prediction on the random test set. **A,B**: Gene-wise correlation between the true and predicted profile, comparing Polarbear with BABEL (A) or with Polarbear co-assay (B), only showing genes that are differentially expressed across cell types. BABEL performance is reported based on the best performing model in each task after a hyperparameter grid search. Each dot is a gene, and numbers indicate the number of dots above and below the diagonal line. P-values are calculated by one-sided Wilcoxon rank-sum test. “Polarbear co-assay” only uses co-assay data to train the model. **C,D**: Peak-wise AUROC, comparing Polarbear with BABEL (C) or with Polarbear co-assay (D). Each dot represents a peak, only peaks differentially expressed across cell types are shown. **E,F**: Gene-wise AUROC. For each marker gene, we calculate the AUROC of its prediction to be higher in cells in corresponding cell type compared to unrelated ones. Polarbear with BABEL (E) or with Polarbear co-assay (F).

Polearbear will be particularly useful if its predictions can be used to derive biological insights, such as predicting genes specific to each cell type. Accordingly, we use a predefined sets of cell-type signature genes that are annotated in the SNARE-seq study [9], as a gold standard, and ask whether Polarbear’s predictions allow us to rediscover these gene to the corresponding cell types. To do so, we calculate the AUROC for a gene’s predicted expression in a one-vs-all fashion for one cell type versus all others. Polarbear is able to predict the gold standard cell type markers correctly with a median AUROC of 0.936. This performance is significantly better than both BABEL (median AUROC = 0.869; Figure 3E) and the co-assay variant of Polarbear (median AUROC = 0.910; Figure 3F).

Polarbear also predicts cell-type specific genes that are not captured based on the true scRNA-seq profiles. Here we focus on the microglia cell type, which consists of only 91 cells in the SNARE-seq dataset. Based on scATAC-seq profiles in the test set, we predict microglia specific genes by calculating AUROC for each gene’s predicted expression in microglia cells against all other cells. Based on Polarbear’s prediction, the *Sall1* gene is specifically expressed in microglia (AUROC = 0.851), but this gene is not highly expressed in microglia based on the observed scRNA-seq profiles (AUROC = 0.498). Interestingly, *Sall1* has been found previously to be a microglia signature gene, and it encodes the transcription factor, Sall1, that maintains microglia identity [25].

### 3.2 Polarbear generalizes to new cell types

Because the amount of available co-assay data is limited relative to single-assay data, a common challenge for models such as Polarbear is to translate between modalities in cell types for which no training data is available. The authors of the BABEL model simulated this scenario by creating a train/test split in which an entire SNARE-seq cell cluster is held out for testing. Accordingly, we also investigate this setting, using BABEL’s train/test split.

First, we investigate whether Polarbear predictions can capture variations within this unseen population. Because the scRNA-seq and scATAC-seq profiles within a cell type are expected to be relatively homogeneous, successfully translating across modalities in this scenario requires the model to capture differences between individual cells, not just differences across cell types. We observe that Polarbear consistently outperforms BABEL and Polarbear co-assay in translation in both directions, suggesting that Polarbear predictions are able to recapitulate meaningful variations across cells within a cell type (Figure 4A-D, Supplementary Figure 1C–D, Table 3).

**Figure 4:**
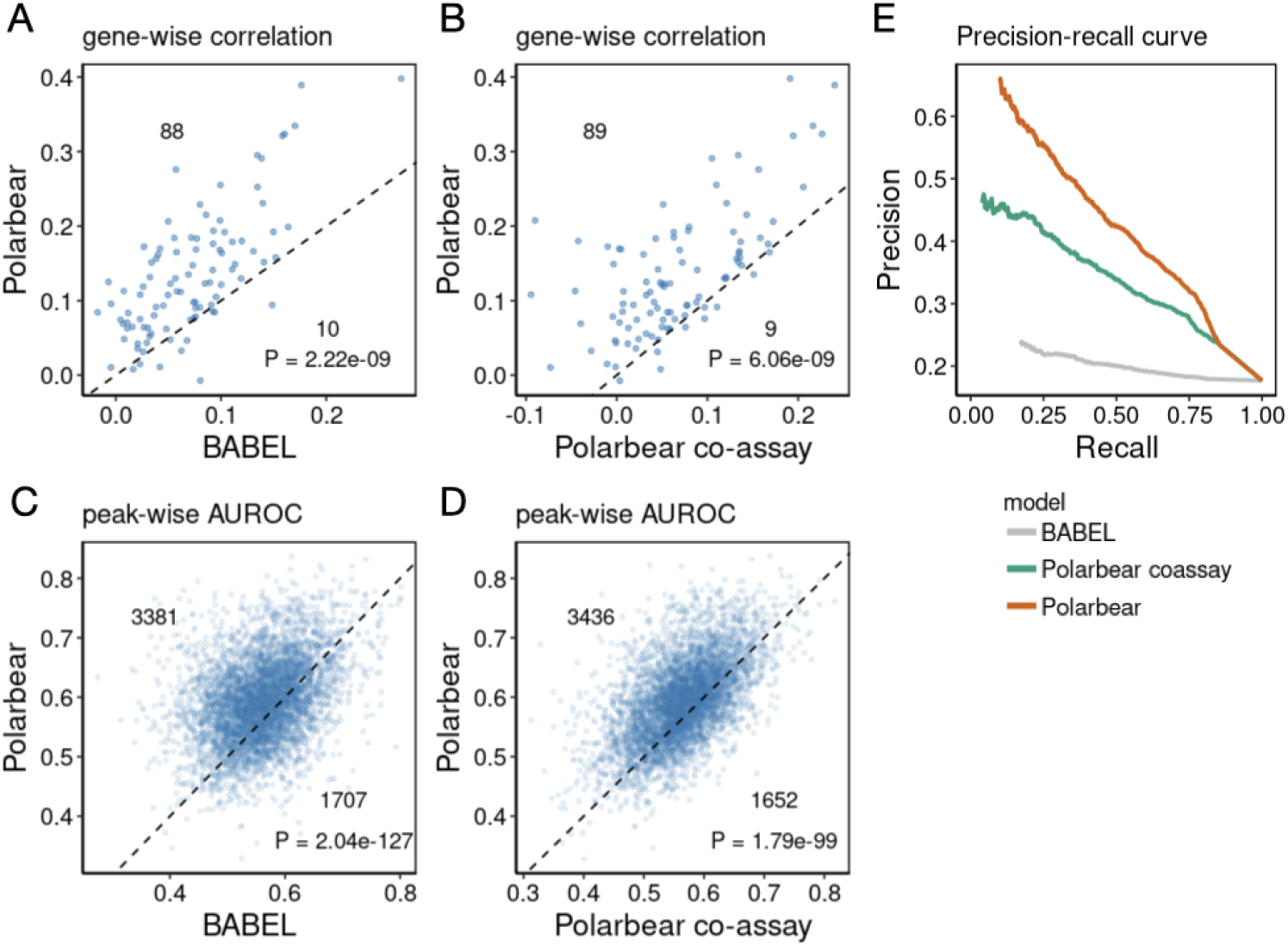
Cross-modality prediction on an unseen cell type. **A,B**: Gene-wise correlation between the true and predicted profile, comparing Polarbear with BABEL (A) or with Polarbear co-assay (B). Each dot is a gene, and numbers indicate the number of dots above and below the diagonal line. P-values are calculated by one-sided Wilcoxon rank-sum test. “Polarbear co-assay” only uses co-assay data to train the model. **C,D**: Peak-wise AUROC, comparing Polarbear with BABEL (C) or with Polarbear co-assay (D). **E**: Precision-recall curve on prioritizing the true set of differentially expressed genes based on differential expression pattern on the predicted profiles.

Next, we test Polarbear’s ability to predict gene expression signatures of the unseen cell population, based only on the scATAC-seq profiles. For this step, we derive the gene signatures for the unseen cell type by identifying genes with significantly higher expression in the unseen cell type compared to cells in the SNARE-seq training set (see Section 2.5) We then calculate a precision-recall curve relative to these cell labels, ranking genes by their differential expression from the predicted profiles. Polarbear’s predictions are able to recapitulate differentially expressed genes, significantly outperforming other methods (Figure 4E). These results suggest that Polarbear can correctly predict intra- and inter-cell type variations, even for cell types for which no co-assay data is available to train the model.

### 3.3 Polarbear can match corresponding cells across modalities

Polarbear can also be used to match corresponding cells from different modalities. Given unpaired single-assay profiles in each modality, we can use Polarbear to match those cells between modalities, supervised by the co-assay data. To simulate this setting, we project the scRNA-seq and scATAC-seq profiles in the held out test set to the bottleneck layer of the Polarbear model, and we match cells from different modalities in a greedy fashion based on Euclidean distance in the latent space. To assess the matching performance, we calculate for each cell the fraction of samples closer than the true match (FOSCTTM) [18]. Polarbear achieves a lower FOSCTTM score than the competing methods in matching cells in the random test set (Figure 5A–B) as well as matching cells within the unseen cell type (Figure 5C–D), suggesting that adding single-assay data improves cross-modality matching.

**Figure 5:**
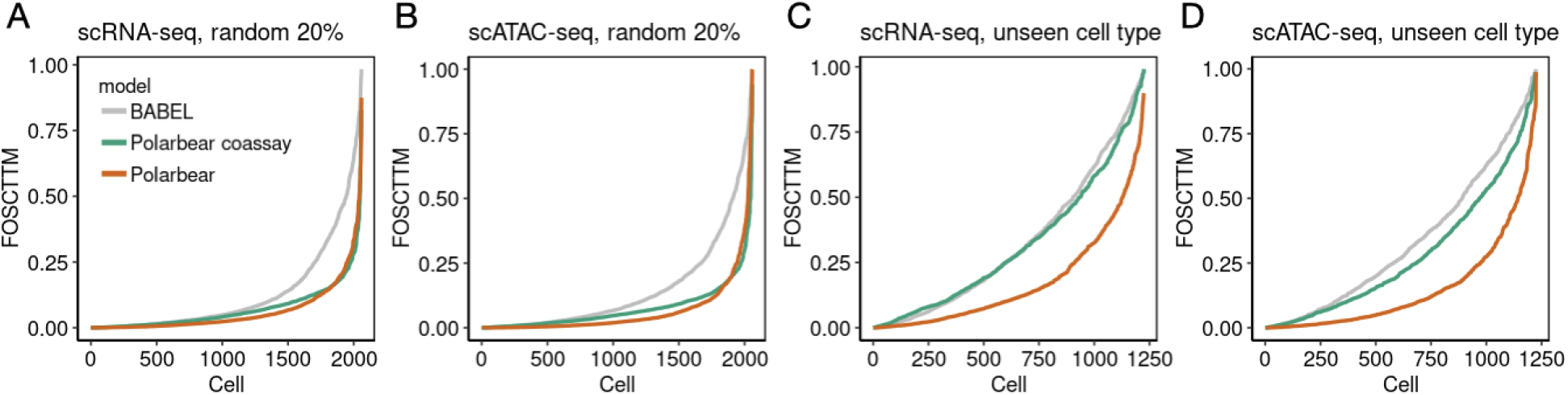
Evaluation of cross-modality matching. The FOSCTTM score for Polarbear (orange), Polarbear trained only with co-assay data (green), as well as BABEL (grey). Cells are sorted based on FOSCTTM score for each method. **A,B**: Matching performance on the random 20% test set, using either scRNA-seq (A) or scATAC-seq (B) as queries. **C,D**: Matching performance on the unseen cell type, using either scRNA-seq (C) or scATAC-seq (D) as queries.

## 4 Discussion

We propose Polarbear, a semi-supervised framework that leverages both co-assay and publicly available single-assay data to translate between scRNA-seq and scATAC-seq profiles. We demonstrate that Polarbear improves upon methods that only train on co-assay data. Polarbear predictions are able to capture cell-type and individual cell-level differences, and can predict missing domain knowledge for cell types without any co-assay data available. Polarbear can be used to generate biological hypotheses in the missing domain, such as inferring differentially expressed genes/peaks between cell types or experimental groups. We expect Polarbear to be used to facilitate biological discoveries on uncharacterized domains at the single-cell level, such as identifying individual cell or subclone specific regulatory elements based on scRNA-seq profiles in tumor samples.

Currently, Polarbear predictions do not improve scATAC-seq predictions as much as scRNA-seq predictions. Possible reasons for this difference are that scATAC-seq profiles are sparse and noisy, and scATAC-seq potentially contains more information than scRNA-seq because a single gene can be regulated by multiple scATAC-seq peaks. We foresee that models taking into account prior knowledge (e.g. DNA-sequence features or regulatory region annotations) may further improve scATAC-seq predictions.

Polarbear can also match single cells across modalities with improved accuracy. We envision that our semi-supervised matching framework could be adapted for aligning the large compendium of publicly available single-assay profiles, so that we can generate new hypothesis (e.g. gene-peak relationships and cell clustering based on joint features) based on the predicted paired scRNA-seq and scATAC-seq profiles.

Thanks to the flexible training framework, the current Polarbear model could, in the future, be combined with pre-trained models from other data domains by learning the translation layer based on a limited number of co-assay data, and thus be generalized to translate between multi-modalities.

---

**Supplementary Figure 1:**
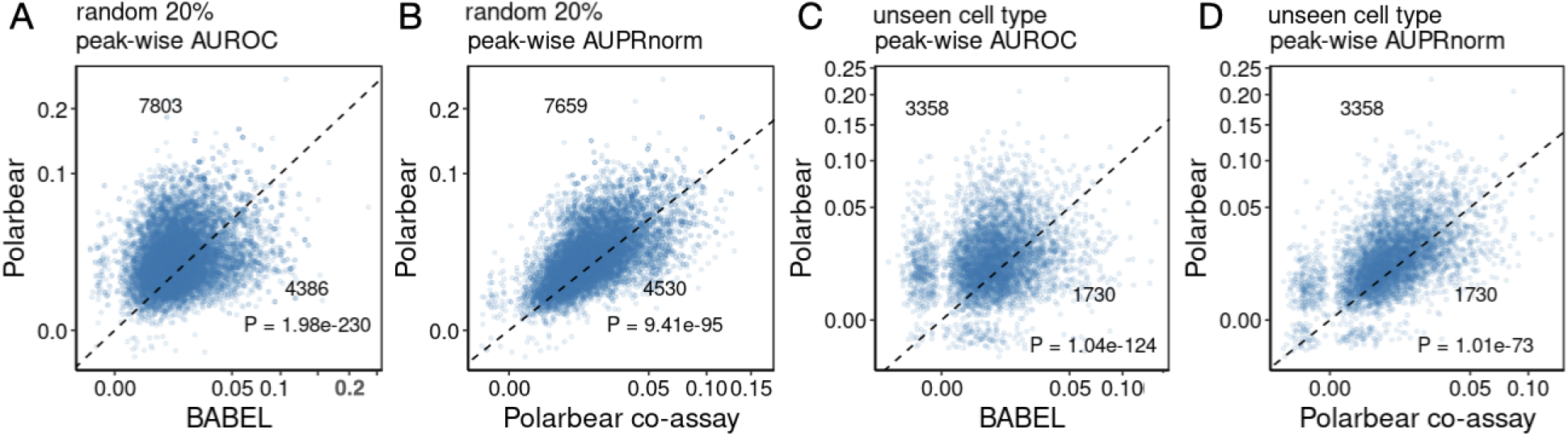
Evaluation of predicted scATAC-seq profile by peak-wise AUPRnorm. **A,B**: Peak-wise AUPRnorm on the random test set, comparing Polarbear with BABEL (A) or with Polarbear co-assay (B). Each dot represents a peak, only peaks differentially expressed across cell types are shown. **C,D**: Gene-wise AUPRnorm on an unseen cell type, comparing Polarbear with BABEL (C) or with Polarbear co-assay (D). Only peaks highly expressed in the unseen cell type compared to other cell types are shown.

## References

[1] A. Gayoso, Z. Steier, R. Lopez, J. Regier, K. L. Nazor, A. Streets, and N. Yosef. Joint probabilistic modeling of single-cell multi-omic data with totalVI. Nature Methods, 18(3):272–282, 2021.

[2] K. E. Wu, K. E. Yost, H. Y. Chang, and J. Zou. Babel enables cross-modality translation between multiomic profiles at single-cell resolution. Proceedings of the National Academy of Sciences, 118(15), 2021.

[3] K. Minoura, K. Abe, H. Nam, H. Nishikawa, and T. Shimamura. A mixture-of-experts deep generative model for integrated analysis of single-cell multiomics data. Cell Reports Methods, page 100071, 2021.

[4] Y. Hao, S. Hao, E. Andersen-Nissen, W. M. Mauck III, S. Zheng, A. Butler, M. J. Lee, A. J. Wilk, C. Darby, M. Zager, et al. Integrated analysis of multimodal single-cell data. Cell, 2021.

[5] M. Lotfollahi, A. Litinetskaya, and F. Theis. Multigrate: single-cell multi-omic data integration.

[6] T. Ashuach, M. I. Gabitto, M. I. Jordan, and N. Yosef. Multivi: deep generative model for the integration of multi-modal data. bioRxiv, 2021.

[7] B. Gong, Y. Zhou, and E. Purdom. Cobolt: Joint analysis of multimodal single-cell sequencing data. bioRxiv, 2021.

[8] J. Cao, D. A. Cusanovich, V. Ramani, D. Aghamirzaie, H. A. Pliner, A. J. Hill, R. M. Daza, J. L. McFaline-Figueroa, J. S. Packer, L. Christiansen, F. J. Steemers, A. C. Adey, C. Trapnell, and J. Shendure. Joint profiling of chromatin accessibility and gene expression in thousands of single cells. Science, 361(6409):1380–1385, 2018.

[9] S. Chen, B. B. Lake, and K. Zhang. High-throughput sequencing of the transcriptome and chromatin accessibility in the same cell. Nature Biotechnology, 37(12):1452–1457, 2019.

[10] C. Zhu, M. Yu, H. Huang, I. Juric, A. Abnousi, R. Hu, J. Lucero, M. M. Behrens, M. Hu, and B. Ren. An ultra high-throughput method for single-cell joint analysis of open chromatin and transcriptome. Nature Structural and Molecular Biology, 26:1063–1070, 2019.

[11] D. Talwar, A. Mongia, D. Sengupta, and A. Majumdar. AutoImpute: Autoencoder based imputation of single-cell RNA-seq data. Scientific Reports, 8:16329, 2018.

[12] T. N. Trong, R. Kramer, J. Mehtonen, G. González, V. Hautamäki, and M. Heinäniemi. SISUA: Semi-supervised generative autoencoder for single cell data. bioRxiv, 2019. https://www.biorxiv.org/content/10.1101/631382v1.abstract.

[13] G. Eraslan, L. M. Simon, M. Mircea, N. S. Mueller, and F. J. Theiss. Single-cell RNA-seq denoising using a deep count autoencoder. Nature Communications, 10:390, 2019.

[14] D. Wang and J. Gu. VASC: Dimension reduction and visualization of single-cell RNA-seq data by deep variational autoencoder. Genomics, Proteomics and Bioinformatics, 16(5):320–331, 2018.

[15] R. Lopez, J. Regier, M. B. Cole, M. I. Jordan, and N. Yosef. Deep generative modeling for single-cell transcriptomics. Nature Methods, 15(12):1053–1058, 2018.

[16] T. Ashuach, D. A. Reidenbach, A. Gayoso, and N. Yosef. PeakVI: A deep generative model for single cell chromatin accessibility analysis. bioRxiv, 2021.

[17] L. Xiong, K. Xu, K. Tian, Y. Shao, L. Tang, G. Gao, M. Zhang, T. Jiang, and Q. Zhang. Scale method for single-cell atac-seq analysis via latent feature extraction. Nature communications, 10(1):1–10, 2019.

[18] J. Liu, Y. Huang, R. Singh, J.-P. Vert, and W. S. Noble. Jointly embedding multiple single-cell omics measurements. In Katharina T. Huber and Dan Gusfield, editors, 19th International Workshop on Algorithms in Bioinformatics (WABI 2019), volume 143 of Leibniz International Proceedings in Informatics (LIPIcs), pages 10:1–10:13, Dagstuhl, Germany, 2019. Schloss Dagstuhl–Leibniz-Zentrum fuer Informatik. PMC8496402.

[19] A. Zeisel, H. Hochgerner, P. Lönnerberg, A. Johnsson, F. Memic, J. van der Zwan, M. Häring, E. Braun, L.E. Borm, G. La Manno, S. Codeluppi, A. Furlan, K. Lee, N. Skene, K. D. Harris, J. Hjerling-Leffler, E. Arenas, P. Ernfors, U. Marklund, and S. Linnarsson. Molecular architecture of the mouse nervous system. Cell, 174(4):999–1014, 2018.

[20] R. Fang, S. Preissl, Y. Li, X. Hou, J. Lucero, X. Wang, A. Motamedi, A. K. Shiau, X. Zhou, F. Xie, et al. Comprehensive analysis of single cell atac-seq data with snapatac. Nature communications, 12(1):1–15, 2021.

[21] Y. E. Li, S. Preissl, X. Hou, Z. Zhang, K. Zhang, R. Fang, Y. Qiu, O. Poirion, B. Li, H. Liu, et al. An atlas of gene regulatory elements in adult mouse cerebrum. BioRxiv, 2020.

[22] J. Harrow, F. Denoeud, A. Frankish, A. Reymond, C.-K. Chen, J. Chrast, J. Lagarde, J. G. R. Gilbert, R. Storey, D. Swarbreck, C. Rossier, C. Ucla, T. Hubbard, S. E. Antonarakis, and R. Guigó. GENCODE: Producing a reference annotation for ENCODE. Genome Biology, 7(Suppl 1):S4, 2006.

[23] A. S. Hinrichs, D. Karolchik, R. Baertsch, G. P. Barber, G. Bejerano, H. Clawson, H. Diekhans, T. S. Furey, R. A. Harte, F. Hsu, et al. The ucsc genome browser database: update 2006. Nucleic acids research, 34(suppl 1):D590–D598, 2006.

[24] A. T. L. Lun, K. Bach, and J. C. Marioni. Pooling across cells to normalize single-cell rna sequencing data with many zero counts. Genome biology, 17(1):75, 2016.

[25] A. Buttgereit, I. Lelios, X. Yu, M. Vrohlings, N. R. Krakoski, E. L. Gautier, R. Nishinakamura, B. Becher, and M. Greter. Sall1 is a transcriptional regulator defining microglia identity and function. Nature immunology, 17(12):1397–1406, 2016.

